# Dynamically arrested condensate fusion creates complex structures with varying material properties

**DOI:** 10.1101/2024.11.15.623768

**Authors:** Nadia A. Erkamp, Ignacio Sanchez-Burgos, Alexandra Zhou, Tommy J. Krug, Seema Qamar, Tomas Sneideris, Ellie Zhang, Kichitaro Nakajima, Anqi Chen, Rosana Collepardo-Guevara, Jan van Hest, Peter St George-Hyslop, David A. Weitz, Jorge R. Espinosa, Tuomas P. J. Knowles

## Abstract

The cell nucleus and cytosol contain numerous biomolecular condensates which dynamically reshape, fuse and split to accomplish precise compartmentalization of the cell material. While it has been observed that some condensates rapidly coalesce, some others only attach to each other, or do not establish persistent interactions over time. Here, we explain these observations through optical tweezers and Molecular Dynamics simulations focusing on two condensate-forming, RNA-binding proteins—FUS and G3BP1—strongly involved in RNA metabolism and stress responses. We find that the fusion of pure droplets formed by these proteins can give rise to multiphase single-component condensates exhibiting notably different densities, architectures, and material properties. Such behaviour is dictated by the relative timescales of condensate fusion and protein internal mixing. A critical parameter controlling this interplay is the extent of ageing that condensates display; e.g., their progressive hardening driven by the accumulation of inter-protein *β*-sheet assemblies over time. Strikingly, different degrees of ageing in fusing droplets can lead single-component condensates to form diverse architectures including concentric drops or two-sided condensates. Overall, our results highlight a mechanism, based on the temporal coupling between ageing, fusion, and mixing rate, by which biomolecular condensates form multiphasic structures with markedly different material properties, and hence potentially distinct biological roles.

## Main

Biomolecular condensates are membraneless organelles which self-assemble via phase separation of biomolecules from a homogeneous solution. Condensates in cells exhibit a wide variety of different functions such as cell signaling [1–3], buffering protein concentrations [4, 5], compartmentalisation [4–7], or genome silencing [8, 9] among others [10, 11]. Some of these condensates can coalesce amongst themselves, like stress granules [12] and some stick together without fusing as for example stress granules with processing bodies [13]. Furthermore, association of condensates can also lead to the formation of multiphasic spherical architectures as the nucleoli [14]. However, some biomolecular assemblies such as paraspeckles and Cajal bodies do not physically attach to each other [15]. The ability of condensates to fuse strongly depends on their composition, post-translational modifications [16, 17] and relative interfacial tensions [18]. Moreover, even when condensates should be able to fuse, such process may be limited by obstacles like the cytoskeleton or the presence of Pickering agents [19–21]. Nevertheless, condensate fusion can also be promoted by active processes, like cytosolic stirring [19].

Whereas condensate fusion has been closely examined as a means-to-an-end to measure material properties [22–25], limited work has been done to understand fusion events involving dynamically arrested condensates similar to those found in cells [26–28]. For example, condensates encountering each other in a cell may have similar composition (symmetric fusion) or may have a different composition (asymmetric fusion) and the impact of this on condensate fusion is not yet understood. Importantly, in cells, arrested fusion has been reported [26], nonetheless, the underlying conditions and pathways leading to such phenomenon remain unclear. The initial condensate composition, protein sequence, or extent of ageing—understood as the degree of kinetic arrest due to inter-protein fibril accumulation—are parameters which can potentially control such behaviour. Lastly, condensate fusion, defined as the emergence of a single spherical condensate, and internal mixing of the material, leading to a homogeneous condensate, are processes that despite being intimately related, may occur at completely different timescales. The interplay between these processes can possibly determine the architecture, shape and material properties of the final assembly. Hence, since condensate viscoelastic properties are strongly related to health and disease [29–31], elucidating how fusion of liquid-like and kinetically arrested condensates takes place is highly urgent to uncover if fusion events are a potential mechanism to further regulate the dynamics of condensates.

In this work, we address these questions through a combination of experimental and computational techniques. We prepare condensates using the proteins FUS (Fused in sarcoma) and G3BP1 (Ras GTPase-activating protein-binding protein 1) (Methods). G3BP1 is a scaffold protein for stress granules, which can recruit FUS and polyadenylated mRNA [32]. Stress granules may play a crucial role in the formation of toxic FUS structures involved with neurodegenerative diseases such as Amyotrophic Lateral Sclerosis (ALS) and Frontotemporal Dementia (FTD) [29, 33]. We prepare the protein condensates with the biocompatible polymer PEG (polyethylene glycol) or polyadenylic acid (poly-rA). We study both FUS-WT and the patient-derived mutant FUS-G156E [34]. Using these condensates, we quantitatively study how symmetric and asymmetric fusion is affected by condensate composition, protein sequence, or extent of ageing. We also quantify the mixing of condensate material during and following a fusion event. We then build on these experimental results with computer simulations, and we focus on the fusion and mixing of FUS low-complexity domain condensates. We identify the conditions that enable protein fusion and assess the extent of ageing at which fusion becomes limited, eventually preventing further mixing. By performing non-equilibrium simulations of symmetric (in which the two interacting condensates are the same) and asymmetric (in which one of the condensates presents more limited diffusion) fusion, we elucidate the influence of protein ageing on condensate fusion. We use our sequence-dependent Mpipi-recharged model [35], which predicts protein phase diagrams and condensates’ viscoelastic properties in near-quantitative agreement with experimental observations [36]. We show that the time it takes for two condensates to fuse is directly related to the extent of ageing— quantified as the concentration of inter-protein *β*-sheets formed within the condensate. Furthermore, our asymmetric simulations align with experimental observations, describing a mechanism by which condensates fuse with limited capacity, forming multiphasic single-component condensates where the material demixes within the resulting droplets. Overall, our experiments and simulations show that fusion and mixing of condensate material may occur on different timescales whose precise coupling is governed by the extent inter-protein *β*-sheet clusters. Therefore, when fusion and mixing become arrested at distinct timescales, this results in the formation of spherical condensates that contain regions with different densities, architectures and viscoelastic behaviour.

## Results and Discussion

### Fusion and mixing of protein condensates with optical tweezers

In this paper, we take condensate “fusion” to mean “obtaining one spherical condensate from two”, while the term “mixing” refers to “obtaining material in which is material from the two original condensates is homogeneously distributed”. We investigate protein condensate fusion and material mixing using an optical tweezers setup (Methods) where the solutions containing the condensates are placed in a airtight well. In Fig. 1, we measure the fusion and mixing time of FUS-WT (wild-type) PEG condensates after 1 hour incubation at 37 °C. Two populations of condensates are prepared, which are fluorescently labeled with different dyes and will be referred to as ‘red’ and ‘blue’ condensates (Methods). Using two optical traps, one for the red and another for the blue condensate, both systems are brought into proximity. By moving forwards the red condensate in trap 1 to the blue condensate in trap 2, fusion is observed (Fig. 1). The force on the two tweezers is measured and confocal fluorescent and brightfield images are taken at the same time. During every second, approximately 1.4 confocal images, 10 brightfield images, and 75000 force measurements in the x and y direction of each trap are recorded.

**Figure 1.**
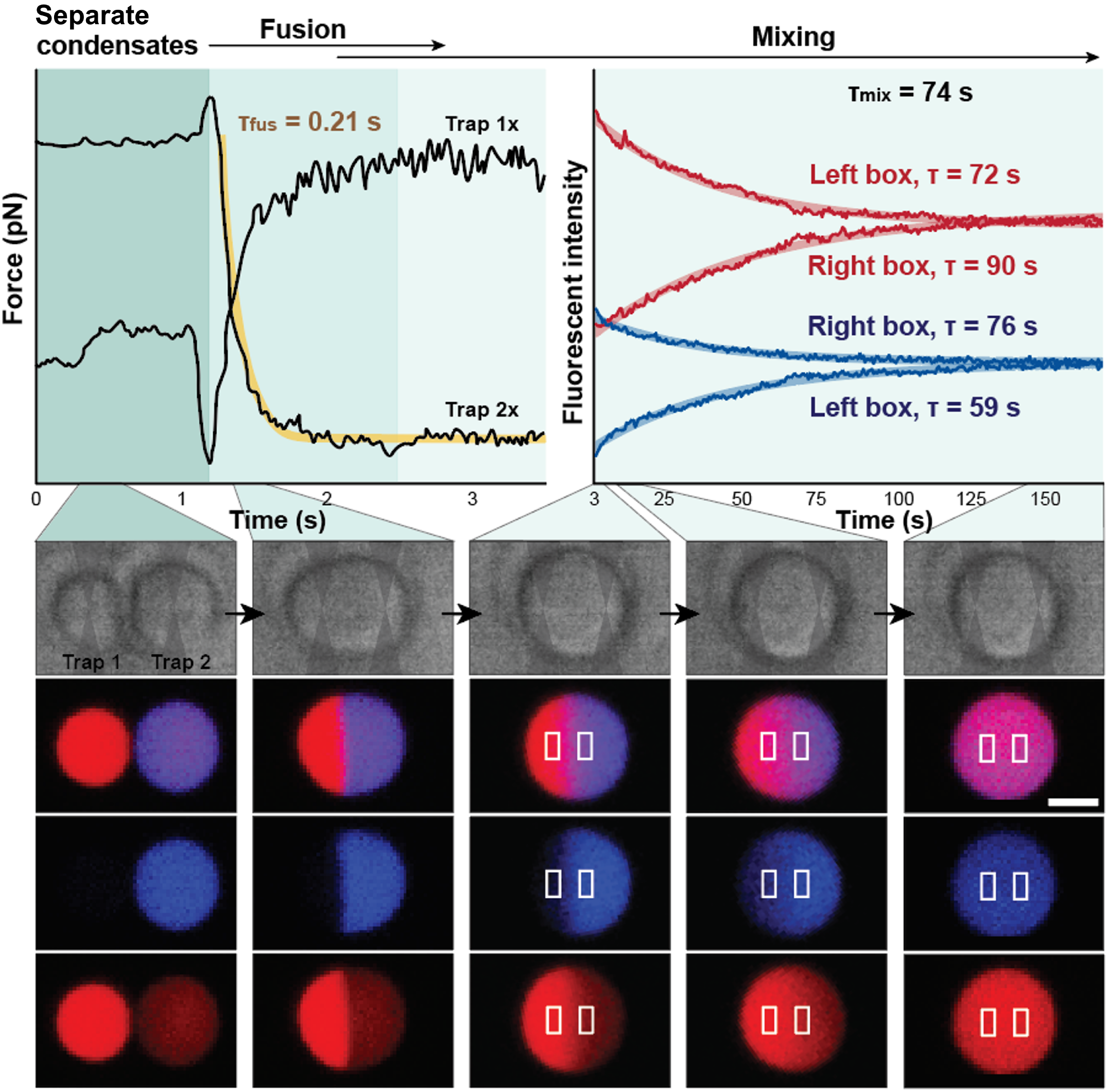
Quantifying the fusion and mixing of FUS condensates. We use a combination of microscopy and optical tweezers to study the fusion of two FUS-WT condensates with different fluorescent labels which are brought in proximity by moving trap 1 towards trap 2. Initial droplets readily fuse into one larger condensate (*τ*_*fusion*_ = 0.21 s). Internal protein mixing requires significantly longer timescales (*τ*_*mix*_ = 74 s). Confocal fluorescent images at the bottom display their state before fusion, during fusion, and 3 frames during mixing. Scale bar represents 3 *µ*m.

The coalescence time for two fusing condensates is obtained using either brightfield and fluorescent images in which the aspect ratio (width over length of the condensates) as a function of time is measured [37]. Notably, we find that the higher frequency force measurements allows us to more accurately determine the fusion time, particularly for fast fusion events. As such, we study the force over time on the tweezers (top left, Fig. 1) where condensates fuse in the x-direction. We note that the force over time experienced by trap 1x is noisier than the signal of trap 2x due to trap 1 being moved during the fusion while trap 2 being held stationary. During fusion, the force on trap 1 increases, while the force on trap 2 decreases. This matches expectations, since trap 1 is pulled at to the right (positive), while trap 2 is pulled towards the left (negative) when the condensates fuse into one. Fusion is finished once the force plateaus to the new value. From the change in the force as a function of time, we determine the fusion time *τ*_*fusion*_. We fit the force over time, *F* (*t*), of trap 2 in the x-direction using:

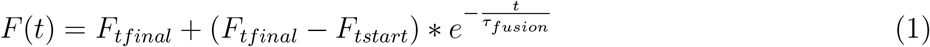

which yields the characteristic fusion time [38, 39], *τ*_*fusion*_ = 0.21 s. The force over time follows the expected trend from Eq. 1 (yellow curve; Fig. 1 top left). Note that the rate at which the force changes is crucial for fitting an accurate *τ*_*fusion*_ instead of the absolute values. Notably, the force that the traps exert on the condensates may influence the time it takes them to fuse [40]. To address this key consideration, we have used a minimum laser power, such that condensates can be seen moving, but not escaping the traps. Additionally, we kept this power constant in all experiments to be able to compare fusion time values. For Newtonian liquids, fusion time can be converted to inverse capillary velocity, the ratio between viscosity, *η*, and interfacial tension *γ*, by normalizing with the average droplet diameter, *d*_*avg*_, before fusion via [40]:

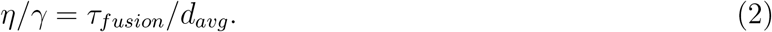

Normalization allows us to quantify the mesoscale fluid properties of the condensates. High interfacial tension drives condensate fusion and indicates condensate stability (further away from critical point), while high viscosity gives a large drag or friction for fusion and lower molecular diffusion [40]. Therefore, elastic-like condensates have a high inverse capillary velocity which may prevent fusion [41].

After fusion, despite observing the formation of a spherical condensate, we realize using confocal images (middle row of bottom, Fig. 1), that our newly formed condensates show a clear boundary between the red and blue material originating from the two starting droplets. This result evidences how fusion occurs at faster rate than internal mixing of the material. Nevertheless, over time, the material does mix and the boundary fades until a mostly homogeneous condensate appears (at 150 seconds; Fig. 1 right bottom panel). As expected for viscous materials [42] with a low Reynolds number, no turbulent mixing is observed. We quantify the mixing of the two initial condensates by tracking the normalized fluorescent intensity of the two wavelengths within the volume of the white boxes depicted in Fig. 1 bottom right; Methods). Since proteins are constantly exchanging between the dense and dilute phase through the interface [43], when potentially accelerates the mixing of the two species via the dilute phase, we select the white-box volumes to be away from the interface to limit such effect. The location of the boxes is at a constant distance from the red-blue starting boundary, and their volumes have constant dimensions, such that values can be compared. We fit the fluorescent intensities over time (top right, Fig. 1) similarly to the fusion time with:

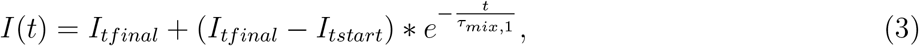

which yields the characteristic mixing time, *τ*_*mix*,1_. Notably, since we track the fluorescent intensity in two locations and for two dyes, we obtain four mixing times (red and blue fitted curves shown in the background of the data; Fig. 1; top right). For each fusion event, we report the average of these values as *τ*_*mix*_. For FUS-WT condensates fusing together, the average mixing time (*τ*_*mix*_) is approximately 74 seconds. Hence, by comparing the fusion and mixing times, we conclude that obtaining a homogeneous condensate (mixing) requires significantly longer timescales—at least 2 orders of magnitude longer—than obtaining a spherical condensate (fusion). We will further explore this difference in detail for asymmetric fusion events (Fig. 3). However, we first explore how symmetric fusion depends on condensate composition, protein sequence, and the extent of droplet ageing.

### Symmetric condensate fusion

Individual biomolecules such as proteins or RNAs can establish an increasing number of interactions of higher strength over time [29, 44, 45]. Terminally, condensates can be viscous Maxwell/Newtonian liquids or elastic Kelvin-Voigt solids [34, 46–48]. Using active processes, like the addition of ATP or other fuel biomolecules, condensates can be driven out-of-equilibrium, creating flow and propulsion among them, and shifting their material properties over time [49, 50]. In the absence of these processes, as in our *in vitro* measurements (Fig. 2), we expect incubation time of condensates at 37 °C to increase the inter-protein *β*-sheet concentration, and hence decrease the inter-molecular mobility over time [28, 51]. Using Eq. 2, we now convert fusion times into the ratio of viscosity by interfacial tension.

**Figure 2.**
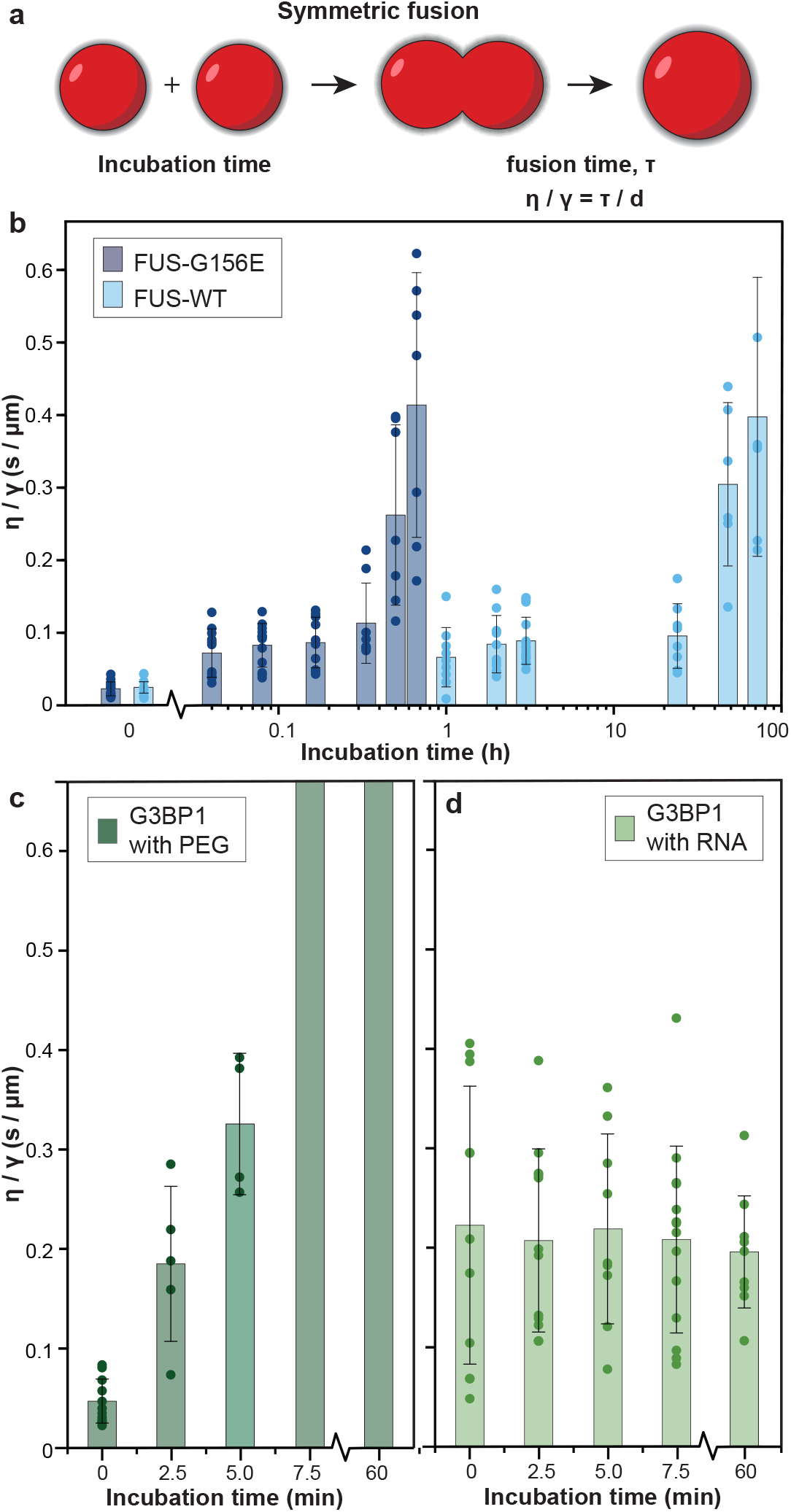
Fusing condensates with similar properties. **a** Symmetric fusion of condensates is performed. Two condensates with similar content, size and age are fused and their *τ*_*fusion*_ is determined. This is normalized using their size to give the ratio of viscosity to interfacial tension. **b** Fusion of FUS condensates incubated for various lengths of time at 37 °C. As FUS-WT condensates (light blue) are stored, the *η/γ* increases over a period of 3 days. Notably, *η/γ* of FUS-G156E slows down to the same extent, except 100 times faster. We can quantify the ageing at high throughput with this setup. **c** Additionally, we study the fusion of G3PB1 PEG condensates, which stop fusing after 7.5 minutes at 37 °C. **d** In contrast, G3BP1 condensates prepared with Poly-rA stay more liquid-like over the duration of the experiment. These results quantify earlier observations (references) and provide a basis for the coming figures.

**Figure 3.**
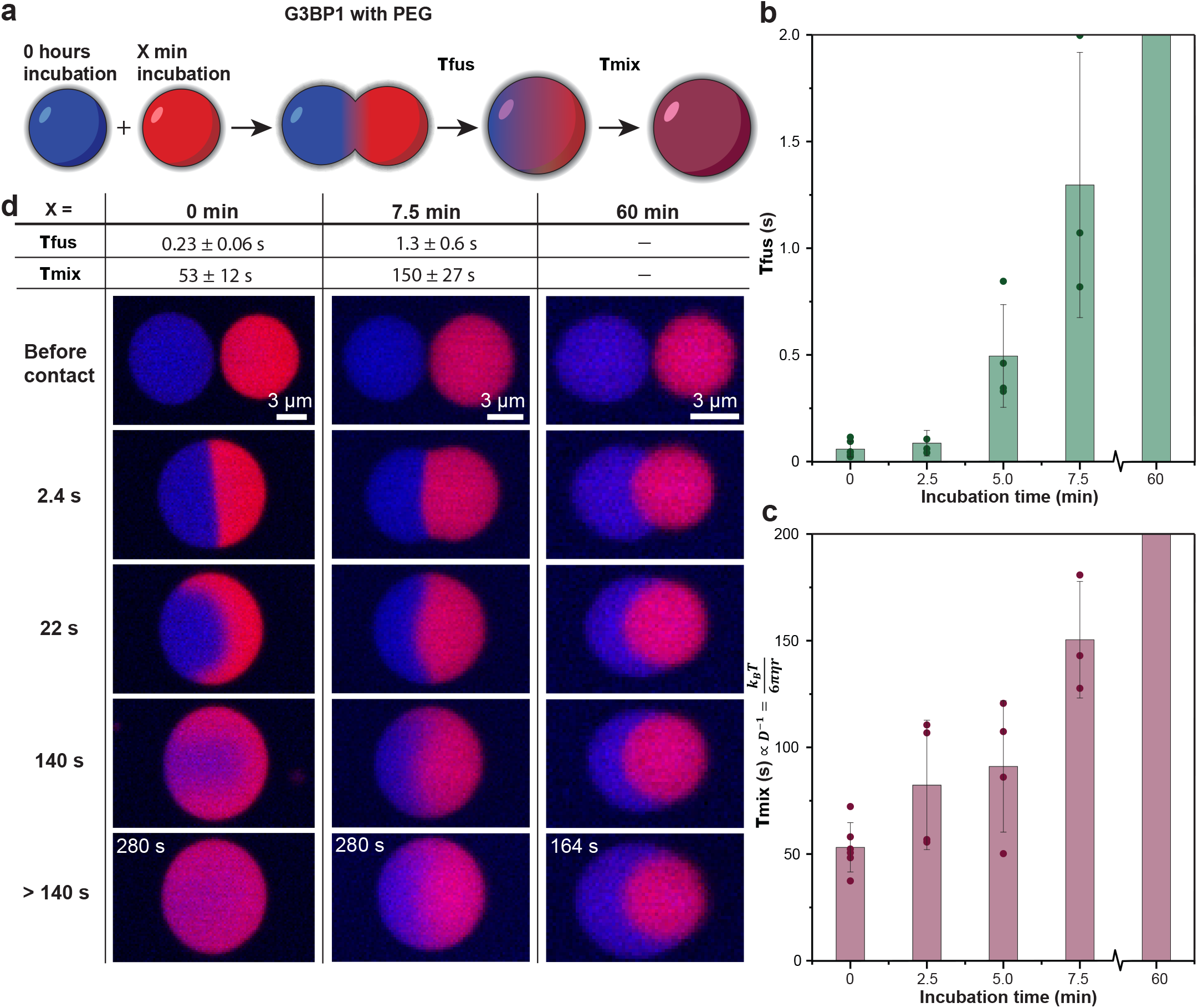
Fusing condensates with dissimilar properties. **a** Asymmetric fusion of condensates is performed. We fuse fresh G3BP1 condensates (blue) with aged G3BP1 condensates (red). **b** Fusion time as function of the age of the blue condensate. At the age of 0 minutes, we have a symmetrical situation similar to Fig. 2c. For the other times, using one aged condensate reduces the fusion time in comparison to two aged condensates. We still observe fusion at 7.5 minutes. **d** Examples of a fusion event with the red condensate aged for 0, 7.5 and 60 minutes. Fusion and **c** mixing times increase with ageing. After 60 minutes ageing, we do not observe a significant amount of fusion or mixing.

First, we focus on the fusion of FUS-WT condensates in presence of PEG. Freshly emerged condensates rapidly coalesce with *τ*_*fusion*_ = 0.068 ± 0.024 s. The use of the high-frequency force measurements enables the accurate determination of such short fusion times. Then, Eq. 2 is used to convert *τ*_*fusion*_ to the ratio between viscosity and interfacial tension (0h, light blue, Fig. 2**b**). Next, condensates are incubated at 37 °C over different time-lapses. We observe a sharp increase on the coalescence time (expressed as *η/γ*). Condensates after an hour fuse in *τ*_*fusion*_ = 0.19 ± 0.13 s, matching our result from Fig. 1, with *η/γ* = 0.063 *s/µm*. As condensates are incubated for longer times, *η/γ* further increases. Such increase confirms our hypothesis of incubation time dramatically decreasing inter-molecular mobility over time and matches earlier work on FUS condensate material properties [34, 52].

To quantify the effect of protein sequence on condensate coalescence, we next study the fusion time of FUS-G156E, a mutant whose condensates have been reported to age significantly faster [51], likely due to cross-*β*-sheet formation [53], although this has not been extensively quantified yet [34]. Without incubation, the fusion time of both FUS sequences is relatively similar (0h, dark blue, Fig. 2**b**). However, the fusion time rapidly rises for the FUS-G156E mutant as a function of the incubation time. After 40 minutes, the inverse capillary velocity is 0.4 ± 0.2 s/*µm*, which is a similar value to that of FUS-WT after 72 hours of incubation. That is, FUS-G156E undergoes condensate ageing approximately 100 times faster than FUS-WT. Interestingly, *η/γ* follows a similar trend for both sequences, except shifted to shorter times by 2 orders of magnitude in the FUS-G156E sequence. Impressively, this mutation of a single amino acid across the 526 residue sequence of FUS dramatically alters condensate maturation. In fact, condensates brought into contact with a longer incubation time than 40 minutes did not fuse into a spherical condensate after 3 minutes, further highlighting the effect of rapid ageing in FUS-G156E.

Next, to further study the effect of condensate composition, we examine the coalescence behaviour of G3BP1 condensates with PEG (Fig. 2**c**) (Methods). G3BP1 PEG condensates age very rapidly at 37 °C (Fig. 2**c**). After 7.5 minutes, condensates stop fusing and only stick to each other forming non-spherical assemblies. We additionally prepare G3BP1 condensates in presence of Poly-rA RNA (Methods). Polyadenylated mRNA is often recruited into G3BP1 scaffolded stress granules [32, 54]. The presence of RNA keeps G3BP1 and other RNA-binding protein condensates significantly more liquid-like [55–57]. For G3BP1 poly-rA condensates, we do not observe a significant change in the inverse capillary force over a period of 1 hour incubation time (Fig. 2**d**). Condensates remain relatively liquid-like and are able to completely fuse. The standard deviation of these measurements in higher than what we have previously obtained. This may be caused by condensates containing slightly different amounts of RNA, which can exchange very slowly once partitioned into the condensates. Matching previous results [55–57], we observe no change in material properties in the absence of condensate aging.

### Interplay of condensate fusion and mixing in asymmetrically aged systems

To further interrogate the role of condensate ageing, we now quantify the fusion and mixing time of G3BP1 PEG condensates with different incubation times. However, to study asymmetric fusion, we give the two condensates a different fluorescent label. Similar to the FUS-WT condensates studied in Fig. 1, the force on the traps is used to measure fusion time and the confocal images are used to monitor the internal mixing time. Fresh G3BP1 PEG condensates (in blue) are brought together with G3BP1 condensates that have undergone ageing for a variable amount of time (red) (Fig. 3**a**). When the ‘aged’ condensate (in red) incubation is 0 minutes, we recover *τ*_*fusion*_ of 0.23 s (Fig. 3**b**) which is consistent with the symmetrical situation. Nonetheless, when aged condensates are incubated for 2.5 or 5 min (Fig. 3**b**) *τ*_*fusion*_ sharply increases, although it remains significantly below than when both condensates are aged (Fig. 2**c**). At 7.5 min, condensates still fuse (Fig. 3**b**), where no fusion was observed when both condensates are aged for this time (Fig. 2**c**). Finally, after 60 min of incubation, fusion is frustrated leading to an aspherical condensate which shows two well defined phases.

For G3BP1 PEG asymmetric fusion, we now analyze the mixing time as a function of the incubation time (Fig. 3**c**,**d**). At 0 min, we observe that the area where the red and blue dense phases are mixing forms a crescent moon-like shape (22s). It appears that it is energetically favorable for the protein labeled with the red fluorescent tag to encapsulate the blue material. This could be caused by the chemical character of the fluorescent dye which induces a small, but noticeable, interfacial tension difference between the two species. The internal mixing time for this system is 53 ± 12 s, which is between 2 and 3 orders of magnitude larger than its fusion time, consistent with our previous findings for FUS-WT in presence of PEG (Fig. 1). As reported for the fusion time dependence with incubation time, the internal mixing time also significantly increases as a function of the ageing time (Fig. 3**c**). At minutes, fusion and mixing becomes significantly slower, but still occurs, as opposed to symmetric fusion (Fig. 2**c**). However, after 60 minutes of incubation, the liquid-like G3BP1 condensate (blue) encapsulates the more solid-like condensate (red), but marginal mixing is observed. We will further discuss these results in combination with computational work connecting condensate fusion and mixing timescales to the emergence of structured cross-*β*-sheets with time.

### Inter-protein *β*-sheet accumulation explains the relation between ageing, fusion and mixing time

We perform non-equilibrium Molecular Dynamics simulations using the Mpipi-Recharged residue-resolution model [58] for the low-complexity domain (LCD) of FUS. The Mpipi-Recharged outperforms the existing residue-resolution force fields for biomolecular phase-separation and drastically improves the description of charge effects in biomolecular condensates, while maintaining the excellent predictions for other systems of its predecessor—Mpipi [35, 36, 59]—and still considering solvation effects implicitly for computational efficiency. We focus on the LCD of FUS (163 residues) for computational efficiency with respect to simulating the full-sequence (526), and also because the LCD is the protein domain where all rich-aromatic segments able to transition into inter-protein *β*-sheets are located [60]. To describe the formation of inter-protein structural transitions in FUS-LCD [60–64], we employ a non-equilibrium algorithm developed by us [27, 57, 65, 66] which considers the intermolecular interaction variation upon the structural transition of low-complexity aromatic-rich kinked segments (LARKS) [60, 67] into cross-*β*-sheets at the atomistic level. This energetic difference between structured cross-*β*-sheets and disordered LARKS is modelled according to atomistic potential-of-mean force calculations recently performed by us for the three FUS LARKS found in its LCD [66].

Our simulations using the Mpipi-Recharged model [58] describe proteins with amino acid resolution (i.e., one bead per amino acid). We prepare systems of different sizes, ranging from ∼200 to ∼600 protein replicas, and allow the condensates to age over time (see Methods for further computational details on the model potential and simulation technical aspects). In Fig. 4**a**, we show the rate at which inter-protein *β*-sheets form as a function of the incubation time in FUS-LCD condensates. With these simulations, we can directly link the incubation time with the concentration of inter-protein *β*-sheets formed, which serves as a direct measurement of the ageing degree. We compute the storage (elastic, *G*^*′*^) and loss (viscous, *G*^*′′*^) moduli as the real and imaginary components of the Fourier transform of *G*(*t*), respectively for FUS-LCD condensates with and without incubation time. In non-aged condensates at low frequencies (*ω*), the elastic response, *G*^*′*^, remains lower than the viscous response, *G*^*′′*^, and the phase angle *δ* = arctan(*G*^*′′*^*/G*^*′*^) is approximately *π/*4. This behavior is characteristic of a liquid-like viscous state where intra- and inter-molecular interactions have sufficient time to relax and dissipate energy. As frequency increases, we observe a crossover point (*G*^*′*^ = *G*^*′′*^) in both aged and non-aged condensates, marking the transition to a solid or glassy state where molecular rearrangements are too slow to dissipate energy and instead store it. In aged condensates, an additional crossover at intermediate frequencies (10^8^ rad · s^−1^) indicates a liquid-to-solid transition, where the storage modulus, *G*^*′*^, surpasses the loss modulus, *G*^*′′*^, once again. Strikingly, the transition from a viscous liquid to a gel-like material occurs at relatively low concentration of inter-protein *β*-sheets; approximately where 20% of the LARKS are engaged in cross-*β*-sheet clusters.

**Figure 4.**
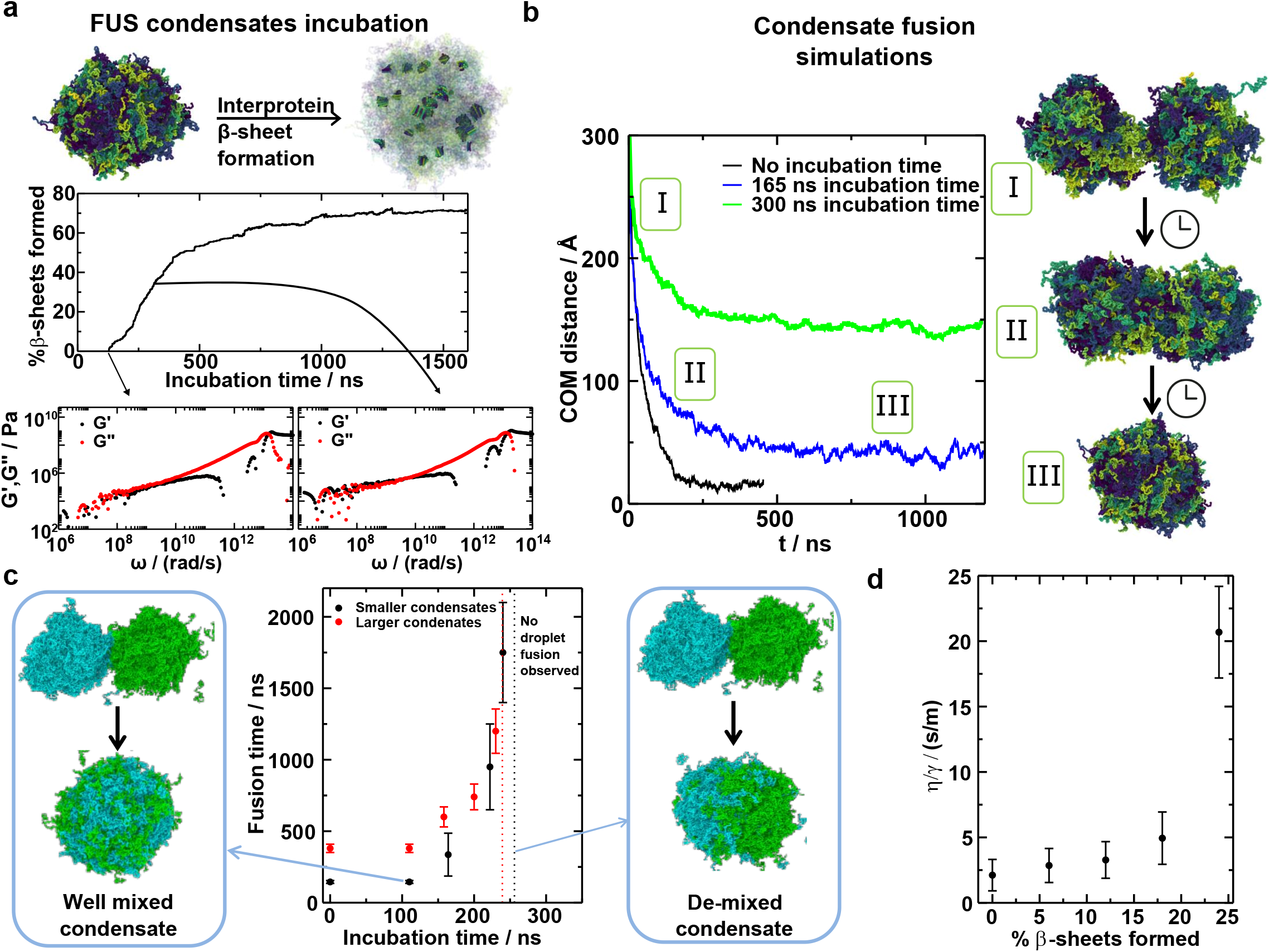
FUS low complexity domain incubation and symmetric fusion simulations. **a** Interprotein *β*-sheet content (expressed in %) as a function of the incubation time. These simulations are carried out allowing the formation of cross-*β*-sheets between different protein replicas. Rendered images showing the FUS LCD condensate as well as the structures formed within are displayed on top of the plot. Material properties change as cross-*β*-sheets are formed. **b** Center-of-mass (COM) distance between the two condensates evolution as a function of time for three different cases as indicated in the legend. The COM distance is computed between the molecules that originally belong to a different condensate. Each curve is the result from averaging four different trajectories with different initial velocities. On the right, images sketching the fusion process of two droplets into a bigger one are shown, where each protein replica is in a different color tone. **c** Fusion time as a function of the incubation time. The fusion time is measured from the fusion simulations plateau (panel **b**) from different extents of ageing. Above ∼250 ns of ageing time (∼22 % of the inter-protein *β*-sheets formed according to panel **a**) we do not observe complete droplet fusion into a well-mixed condensate (illustrated on the left) but rather a de-mixed condensate (as shown on the right). These simulations are performed for two different system sizes, as indicated in the legend, where in the smaller condensates system, each droplet contains ∼200 protein replicas, while the large condensates contains ∼600 replicas each.

We perform condensate fusion simulations by preparing configurations in which two spherical droplets are initially in contact, and can undergo fusion into a single larger condensate (Fig. 4**b**). To assess the time to fuse, we track as a function of time the center-of-mass (COM) distance between the molecules that initially belong to separate condensates. In Fig. 4**b** we plot the COM distance as a function of time for three different cases of symmetric fusion that differ in the droplet incubation time. For the case with no incubation time (black curve), the COM distance rapidly decays to values below ∼25 Å, which approximately corresponds to the value of the radius of gyration (*R*_*g*_) of a single FUS-LCD protein [68–70]). As such, a COM distance of 25 Å is considered as the threshold for achieving complete fusion. For the intermediate case with 165 ns of incubation time (blue curve), the system also evolves into a state in which the COM distance is below the size of a single-protein *R*_*g*_. However, fusion takes considerably longer (near 1 microsecond) than for condensates with no incubation time. Lastly, condensates with longer incubation times (300 ns; green curve) fail to completely fuse, as indicated by the COM distance between molecules of each droplet remaining larger than 100 Å. Importantly, we note that since our protein model has implicit solvent to be computationally efficient, the dynamics are ∼3 orders magnitude artificially faster than experimental measurements. Nevertheless, the predicted trends upon rescaling have been shown to successfully reproduce the relative changes in viscoelastic properties that condensates formed by different mutated sequences display [36].

In Fig. 4**c**, we show the fusion time as a function of the incubation time for FUS-LCD condensates. Symmetric fusion time increases exponentially with the concentration of inter-protein *β*-sheets. Such trend is fully consistent to the optical tweezers measurements from Figure 2**b**. Moreover, these results reassure the concept that the material properties of biomolecular condensates are time-dependent, and can progressively drift from initial liquid-like behaviour to gel and solid-like states with increasing difficulty to flow and coalesce [51]. When short or no incubation time has elapsed (and few inter-protein *β*-sheets are formed), droplets completely coalesce, as illustrated in Figure 4**c**(left). However, above a certain incubation time (in our model ∼250 ns) and concentration of inter-protein *β*-sheets (∼20 % of the total possible cross-*β*-sheets that FUS-LCD can establish), we observe that condensates are no longer able to fuse (at least within feasible computational timescales, vertical dotted line in Figure 4**c**). This leads to condensates in which proteins initially coming from a different droplet remain in the same region, even though the two droplets reshape to form a larger almost spherical condensate (Figure 4**c**, right panel). Such partial reshape of the condensate into a semi-spherical droplet is driven by the surface tension, which tends to minimize the area exposed between the coexisting (condensed and diluted) phases. This is alike to the confocal images after 60 min of incubation time in Figure 3**d** for G3BP1 PEG condensates. We also interrogate the effect that condensate size has on fusion time (Fig. 4**c**). As expected, larger condensates (of 600 protein replicas each) take longer times to reach a completely fused and mixed state than droplets of 200 proteins. Nonetheless, the limit of incubation time to achieve complete fusion remains almost the same, highlighting the fact that longer incubation times affect the viscoelastic properties of all condensates in a similar manner, regardless of their size. To fully compare these results to those of Fig. 2, we evaluate how *η/γ* evolves as a function of the percentage of cross-*β*-sheets formed in the condensates (4**d**) (for details on *η* and *γ* calculations see Methods section). A sharp increase in *η/γ* is observed at 25 % of cross-*β*-sheets formed, which matches with the incubation time at which condensates remain attached, but internally demixed. Such behaviour is consistent with the experimental trend of *η/γ* in FUS-WT condensates symmetric fusion as a function of the incubation time (Fig. 2).

### Fusion of droplets with varying degrees of ageing can induce single-component multiphasic condensates

In addition to our symmetric fusion simulations, we prepare configurations in which one of the two condensates has undergone ageing, while the other had no incubation time. Once the condensates are in contact (Fig. 5**a** and 5**b**), the evolution of fusion is tracked by computing the COM distance of the proteins belonging to each initial condensate (as performed in Figure 4**b**). Since in asymmetric fusion the proteins of the non-aged condensate diffuse faster than those of the incubated one, fusion times can be computed through the COM distance parameter, but mixing times require a further analysis of the protein probability distribution of both species across the newly formed condensate. By computing the radial density profile from the center of the newly formed condensate, we can monitor the degree of internal mixing at different time windows (Figs. 5**c** and 5**d**).

**Figure 5.**
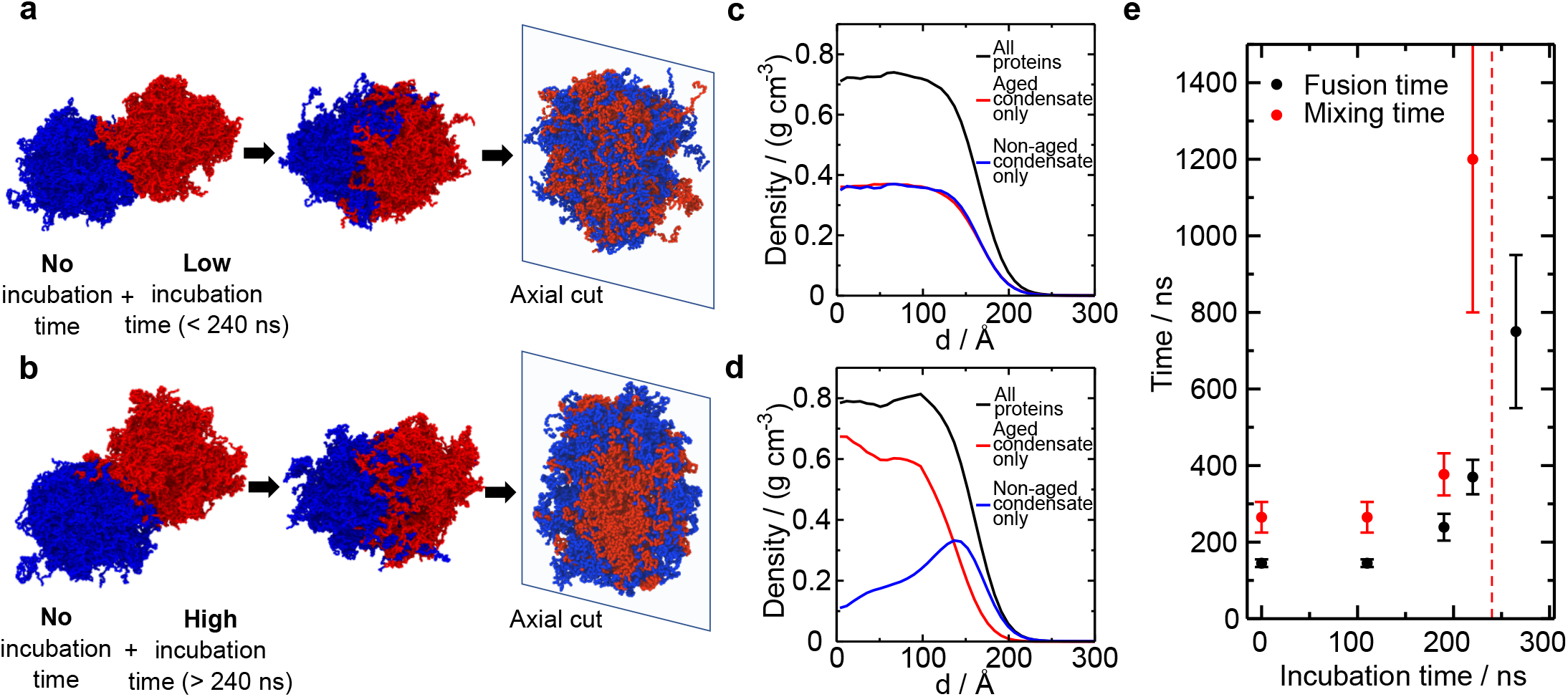
FUS low complexity domain asymmetric fusion simulations. **a**,**b** Rendered images sketching the asymmetric fusion time evolution. The molecules coloured in blue depict the molecules from a condensate with no incubation time, while the red one correspond to the molecules coming from a pre-aged condensate. In **a** we illustrate the case in which there is a low content of inter-protein *β*-sheets formed in one of the condensates, while in **b** the aged condensate has a high content of inter-protein *β*-sheets. In the last image of both panels we show an axial cut of the formed condensate to illustrate the molecular arrangement. **c**,**d** Radial density profiles, calculated from the center of the condensate towards the interface, differentiating between the molecules coming from the aged and non-aged condensate. In **c** we show the profile for the low incubation time case, while in **d**, the incubation time for the aged condensate is higher. **e** Fusion and mixing time in asymmetic fusion simulations. The fusion time is measured as the time it takes to the COM distance below the size of a protein replica, while the mixing time is measured as the time at which the aged and non-aged radial density profiles become equivalent. The vertical dashed line indicates the maximum incubation at which we are able to observe good condensate mixing in asymmetric fusion simulations.

We investigate asymmetric fusion for two different cases: (1) in which the aged condensate was incubated for a short period of time, leading to a concentration of cross-*β*-sheets lower than 20% (Fig. 5**a**); and (2) for a condensate in which the incubation time was longer, and over 20% of cross-*β*-sheets were formed (Fig. 5**b**). As in G3BP1 PEG condensates (Fig. 3), short incubation times for the aged droplet enables fusion and mixing of their material—i.e., the radial density profile eventually shows equivalent density probabilities for both species in the fused condensate (Fig. 5**c**). However, when the concentration of inter-protein *β*-sheet structures increases beyond a certain point, which for FUS-LCD is approximately 20%, protein self-diffusion across the aged condensate dramatically decreases (being G’ *>* G” at high frequencies; Fig. 4a), and the mixing of the two condensates becomes restrained. In this case, the formation of a two-phase concentric condensate with a higher density of the aged phase at the inner core is found (Figure 5**b**). The proteins from the non-aged condensate encapsulate the aged inner phase forming a two-phase demixed droplet that exhibits spherical shape to minimize its exposed area to the protein diluted phase, therefore minimizing its surface tension [71, 72]. Such behaviour for FUS-LCD is fully consistent with the experimental observations for G3BP1 condensates after 60 minutes of incubation as shown in Figure 3**d**).

Furthermore, we measure the mixing time of the asymmetric fusion simulations by monitoring the time for the radial distribution probability of both species (aged and non-aged droplets) to become equivalent (such as that shown in Fig. 5**c**). The fusion and mixing times for FUS-LCD droplets as a function of the incubation time of the aged condensate are shown in Fig. 5**e**. As in Figs. 3**b-c** for G3BP1, both curves exponentially increase with the incubation time of the aged droplet. Above ∼240 ns of incubation (which corresponds to ∼20% concentration of cross-*β*-sheets), we no longer observe protein internal mixing between the condensates. Nonetheless, we are still capable of measuring fusion times above such limit since condensates that are effectively demixed in a two-layer architecture (Fig. 5**b** and **d**), can still form spherical droplets. Consistent with our experimental observations, our simulations from Fig. 5**e** indicate that mixing times are substantially longer than fusion times. Remarkably, our results suggest that the limit for cross-*β*-sheet structures to hinder protein mobility within a condensate is relatively low, with only ∼20% of the FUS-LCD LARKS engaging into inter-protein *β*-sheet structures, while still being sufficient to kinetically arrest the whole system. Hence, the progressive accumulation of highly stable cross-*β*-sheets over time can give rise to the emergence of multiphasic condensate architectures with different shapes (e.g., concentric (Fig. 5**d**) or two-side Fig. 3**d**), densities, and material properties through a fusion mechanism. Moreover, we note that in our simulations for FUS-LCD, the mixing time is only 2-4 times larger than the fusion time, while in G3BP1 PEG experiments, the ratio of mixing over fusion time can be up to two orders of magnitude. We ascribe such significant difference to two important factors: (1) the much shorter protein length of FUS-LCD (163 residues) compared to G3BP1 (932 amino acids for the protein dimer) which drastically increases its protein diffusion coefficient [73, 74]; and (2) the droplet size of simulations vs. experiments which substantially modifies the condensate surface to volume ratio, and thus, the fusion vs. internal mixing times. We also find—in agreement with G3BP1 fusion experiments—that asymmetric fusion of FUS-LCD aged droplets (Fig. 5**e**) occurs significantly faster than symmetric fusion of aged condensates (Fig. 3**c**), showing that just below the critical cross-*β*-sheet concentration (e.g., 200 ns) asymmetrical fusion is ∼3 times faster than symmetrical fusion for the same incubation time of the aged condensates.

## Concluding remarks

In a cell, two condensates encountering each other can coalesce into one, but their fusion can also be dynamically arrested [26, 27]. The extent to which condensates are dynamic is strongly related to various diseases [29–31]. In this work, we explored the conditions and mechanisms regulating the fusion and mixing of protein condensates, with particular focus on how ageing impacts these processes. Using optical tweezers and high-frequency force measurements, we observed that fusion of protein condensates into a spherical droplet occurs rapidly, yet internal mixing of their components is comparatively much slower, requiring several orders of magnitude longer to complete. This difference suggests that while condensates may easily coalesce into a single entity, the material within them may remain compositionally separated for long periods. As observed in the fusion of FUS-WT and G3BP1 condensates, the fusion time, and the viscosity-to-interfacial-tension ratio (*η/γ*) increases as condensates age, with aged condensates sometimes failing to fuse completely, instead forming non-spherical, phase-separated structures. This aging-induced rigidity is even more pronounced in mutant variants like FUS-G156E, which demonstrated accelerated aging, leading to inhibited fusion and mixing within a much shorter time frame compared to the wild-type (Figure 2).

Our observations in asymmetric fusion experiments further highlight the effects of aging, where non-aged condensates were paired with aged ones. When these dissimilar condensates fused, a notable core-shell structure formed, in which the more aged, gel-like condensate constituted the core, surrounded by a fresher, liquid-like shell (Figure 3). This structure not only prevents full internal mixing but also suggests a mechanism for aged condensates to maintain their composition without influencing the surrounding material. This architecture may therefore play a protective role, containing the spread of ageing effects and potentially shielding cellular environments from further aggregation or phase separation that could lead to dysfunction. These diverse architectures arise based on the timing of fusion, rather than only based the molecules themselves. Given the importance of structure for condensate function [75], these observations are relevant for understanding normal and diseased cell function. Our Molecular Dynamics simulations support this by showing that beyond a certain threshold of inter-protein *β*-sheet accumulation, protein condensates are no longer able to fully fuse (Figure 4). Instead, they form stable, layered structures with clear phase boundaries, which also minimize the interfacial free energy between the protein condensed and diluted phases (Figure 5).

In summary, this study advances our understanding of the factors influencing protein condensate fusion and mixing, highlighting the role of ageing in these processes. Dynamically arrested fusion occurs when both or one condensates have exceeded a critical beta-sheet content, leading to condensates sticking or a core-shell structure respectively. Spherical condensates without clear internal interfaces can contain regions with different compositions and/or materials properties.

## Methods

### Materials

The Sf9 cells, AlexaFluor 488/647 NHS ester (succinimidyl ester) were obtained from ThermoFisher Scientific. PEG(20.000), Poly-rA, CAPS (N-cyclohexyl-3-aminopropanesulfonic acid), KCl, glycerol, DTT (Dithiothreitol) and EDTA (ethylenediaminetetraacetic acid) was purchased from Sigma Aldrich. The pACEBac2 vector was purchased from Geneva Biotech. Ni-NTA affinity resin was obtained from Protein Ark. NiAmylose affinity, TEV protease was obtained for New England Biolabs. Protease inhibitor cocktail ordered and used as prescribed by Roche. HALO-AF488 and HALO-AF647 were purchased from Promega.

### Expression and purification of FUS-WT and FUS-G156E

Sequences encoding for his-tag, FUS residues 1 to 466 (with and without G156E mutation) with HALO and MBP tag were placed into pACEBac2 vectors using Gibson cloning. Similarly, we obtained the pACEBac2 vector for his-tag, G3BP1 residues 1 to 526, TEV protease cleaving site and MBP tag. Sf9 cells were infected with baculovirus to express the proteins. Three days after infection, cells were pelleted by spinning at 4000 rpm for 30 minutes. Pellets for purifying FUS were resuspended in 0.1% CHAPS, 1M KCl, 5% glycerol, 1 mM DTT, protease inhibitor cocktail and 50 mM Tris at pH=7.5. In the case of G3BP1, we used 0.1% CHAPS, 1M KCl, 2 mM EDTA, 1 mM DTT, protease inhibitor cocktail and 50 mM Tris at pH=7.5. Proteins were purified over Ni-NTA affinity column and amylose affinity column. Excess tags were cleaved off using TEV protease. Optionally, to FUS, HALO-AF488 or HALO-AF647 was added and optionally to G3BP1, AF488 and AF 647 NHS ester was added. After overnight incubation, size exclusion chromatography was performed in the same buffers as previously, except no CAPS, glycerol or protease inhibitor was added. Fluorescently labelled material was mixed with unlabelled material to obtain 5 mol-% labelled material.

### Optical tweezer wells

Slabs of PDMS (Corning) of 3 mm in height were prepared by mixing the PDMS base and crosslinking agent in a 10:1 ratio and baking at 65 °C for 1 hour. Wells were punched in the PDMS using a 6mm Harris Uni-Core puncher, after which the PDMS was cleaned by sonication in ethanol and then baked at 65 °C for 30 min. The PDMS slabs and 1.5 cover glasses (24×60 mm, DWK life sciences) were activated in an oxygen plasma oven (30s, 60% power, Femto, Diener Electronics) and then bonded together. Samples are placed in the well and covered by a cover glass slide (18×18 mm, Academy).

### Samples for optical tweezer experiments

All samples are prepared in a buffer of 150 mM KCl, 1 mM DTT and 50 mM Tris pH = 7.5. When studying only fusion time, a typical sample is 20*µ*L in volume and contains 5 *µ*M protein (5-molar% fluorescently labelled) and 5% 20k PEG or 10 ng/*µ*L Poly-rA in buffer (figure 2). When mixing time is being studied as well (figure 1, 3, 4), we require condensates with different fluorescent labels (AF488/647 NHS ester). Two solutions of 10 *µ*L containing each 5 *µ*M protein (5-molar% fluorescently labelled) and 5% PEG or 10 ng/*µ*L Poly-rA in buffer are prepared and then placed on different sides of the well. 5 *µ*L of buffer (with 5% PEG if PEG is present in the condensate samples) is placed in between to prevent rapid mixing.

### Optical tweezer setup

A Lumicks C-trap instrument with brightfield imaging, 488 and 637 nm fluorescent confocal microscopy, and dual-trap capabilities was used. Condensates are moved towards each other and held in position at 0.5% power of the trapping laser at 1024nm, to prevent excessive heating of the sample. At this power, condensate can be seen diffusing small distances around the focus of the trapping laser. To fuse condensates together, they are brought in proximity by moving the trapping lasers, at which point we wait until they come into contact due to their movement. A brightfield video, confocal video and relative changes in the force on the two traps in the x and y direction are recorded. The *τ*_*fusion*_ depends on the viscosity and surface tension of the condensates, but also on their size [38, 39]. To reduce the effect of condensate size on this number, we have studied condensates of similar sizes throughout this manuscript.

### Fusion and mixing time analysis

Jupyter notebook is used to extract an .csv file with the force in 1x, 1y, 2x and 2y as a function of time from the .h5 files as output by the Lumicks C-trap software. We obtain approximately 75000 datapoints per second per direction and trap combination. If the fusion took place in the x direction, we fit either 1x or 2x, depending or which trap was not recently moved. Similarly, we use either 1y or 2y if the fusion occurred mostly in the y direction. We use Matlab and the Matlab-provided app “curve fitting toolbox” to fit the force as a function of time according to formula 1 (main text). Mixing time is measured using formula 2 using the same curve fitter. Notably, the reported *τ*_*mixing*_ is the average of 4 fitted curves, while the fusion time is based on fitting only one curve.

### Software

The .h5 files of the confocal data are exported to .tiff files using the Lakeview software. The confocal data is further analysed to obtain the mixing time using Fiji (2.3.051). Figures were prepared with Origin (2017), Adobe Illustrator (27.8.1).

### Computational details

Our Molecular Dynamics simulations are performed in the NVT ensemble (i.e. constant number of particles (N), volume (V) and temperature (T)), for which we use a Nosé–Hoover thermostat [76, 77] with a relaxation time of 5 ps, in order to keep the temperature constant at 273 K (being 300 K the critical temperature for phase separation for the FUS LCD [58]). We perform all our simulations using the LAMMPS Molecular Dynamics package [78]. Periodic boundary conditions are used in the three directions of space. The integration time step chosen is 10 fs. We use the coarse-grained amino acid level resolution model Mpipi-Recharged [58]. For further details on the potential equations please see Ref. [58]. In all our simulations, we keep the salt concentration at 150 mM.

We perform our incubation simulations allowing the formation of inter-protein *β*-sheets to take place, according to the scheme developed in Refs. [27, 65, 66]. We first identified the regions of the FUS sequence that are prone to forming inter-protein secondary structures, as reported in reference [60]. These are low-complexity aromatic-rich kinked segments (LARKS), and in the FUS LCD sequence we can find the following: _37_SYSGYS_42_ (PDB code 6BWZ), _54_SYSSYGQS_61_ (PDB code 6BXV), and _77_STGGYG_82_ (PDB code 6BZP) [60]. In Ref. [66] the potential of mean force of the structured and disordered conformations of such regions was calculated, indicating us what the interaction difference between before and after such structure is formed is. Our incubation simulations consist of non-equilibrium Molecular Dynamics runs in which, when the aforementioned regions are found in space, an effective change in the pairwise interaction of the involved residues (according to the PMF calculations) is performed. We use a distance criterion, in which at leas the central residues of four LARKS of the same kind must meet within a cutoff distance of 14 Å. This distance is chosen so that the formation of the inter-protein *β*-sheets takes place within an accessible timescale. Moreover, the requirement of at least four LARKS of different protein replicas meeting simultaneously is set so that the structures reported in Ref. [60] can be sustained. In this way we model the formation of inter-protein *β*-sheets. This is done with the *fix bond/react* command [79] available in LAMMPS.

In our condensate fusion simulations, two configurations of spherical condensates are placed in direct contact, so that the fusion time can be readily measured. During the preparation of the initial configuration, any protein replica accidentally overlapping in space with another is removed. In this simulations, the amount of inter-protein *β*-sheets is fixed i.e. the condensates cannot form new inter-protein structures during this simulation, in order to measure the fusion time after a fixed incubation or ageing time.

We measure the viscosity (*η*) from independent bulk simulations performed at the equilibrium density of the protein condensed phase in the canonical ensemble for different degrees of ageing. In these simulations, we first obtain G(t) from the six independent components of the pressure tensor [52] and then, using the Green-Kubo relation, integrate G(t) to obtain *η*:

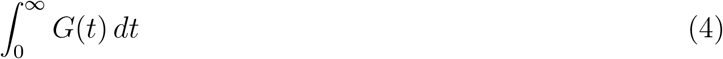

For this integration, we use the Reptate toolkit [80] because, while the G(t) is smooth at short timescales, for long timescales it is required to fit G(t) to a series of Maxwell modes (*G*_*i*_exp(−*t/τ*_*i*_))) equidistant in logarithmic time, to then integrate analytically the region of long timescales. For further details on this method see Ref. [52].

Moreover, from additional Direct Coexistence simulations we measure the interfacial free energy (*γ*) of the condensate by using the Kirkwood-Buff expression [81]:

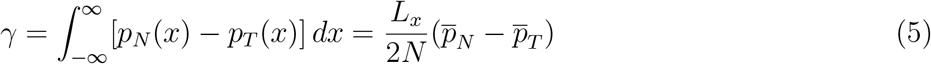

where *L*_*x*_ is the long side of the simulation box (perpendicular to the surface section in our simulations), N is the number of droplets in our system, and *p*_*N*_ and *p*_*T*_ are the normal an tangential components of the pressure tensor with respect to the surface of the system (note that the tangential component is averaged over two tangential directions). We estimate the droplet surface tension with the capillary approximation as the *γ* at planar interface.

## Data Availability

All data generated or analysed during this study are either included in this article and its Extended Data Figures or are available upon request by contacting the corresponding author.

## Acknowledgements

We thank Samuel Cohen, Mina Farag and Rohit V. Pappu for discussions. We thank Luay Joudeh, Katherine Stott, Roman Renger, Emma Verver and Ann Mukhortava for support with the use of the C-trap optical tweezer machine. This work was funded by an ICMS fellowship (N.A.E.), the Royall Scholarship (N.A.E.), the UKRI EPSRC under the UK Government’s guarantee scheme EP/Z002028/1 (I.S.-B., R.C.G.), the European Union’s Horizon 2020 research and innovation program under the Marie Skłodowska-Curie grant MicroREvolution (agreement no. 101023060; T.S.) the Newman Foundation (T.P.J.K., T.S.), Canadian Institutes of Health Research (Foundation Grant and Canadian Consortium on Neurodegeneration in Aging Grant, P. St G.H.), Wellcome Trust Collaborative Award 203249/Z/16/Z (P. St G.H., T.P.J.K.), the US Alzheimer Society Zenith Grant ZEN-18-529769 (P. St G.H.), the European Research Council under the European Union’s Horizon 2020 research and innovation program through the ERC grant DiProPhys (agreement ID 101001615;T.P.J.K.) and the Newman Foundation (T.P.J.K.).

## Author Contributions

Should be updated N.A.E., T.S. and T.P.J.K. conceived the study. N.A.E., I.S.-B.,A.Z., T.J.K., S.Q., T.S., D.Q., E.Z., K.N., A.S., J.H., P.G.-H., D.A.W., J.R.E. and T.P.J.K. performed investigations and interpreted the results. P.G.-H., D.W. and T.P.J.K. acquired funding. N.A.E., I.S.-B., J.R.E. and T.P.J.K. wrote the original drafts, all authors reviewed and edited the paper.

## Additional information

Data generated during the study are available on request from the corresponding authors.

